# Spatial analysis of environmental factors at found locations of orphaned mammals (*Didelphis virginiana*, *Procyon lotor*, *Sciurus carolinensis*, and *Sylvilagus floridanus*) in Champaign County, Illinois, USA

**DOI:** 10.1101/2023.07.20.549871

**Authors:** Stephanie Heniff, William Sander, Colleen Elzinga, Csaba Varga, William Marshall Brown, Samantha J. Sander

## Abstract

Interactions between people and wildlife are increasing as developments encroach on nature. Concurrently, neonatal and juvenile mammals are presented to rehabilitation centers as real or perceived “orphans”, believed to be lacking appropriate parental care for survival. Four common orphaned mammals presented to wildlife rehabilitation facilities are the Virginia opossum (*Didelphis virginiana,* DIVI), common raccoon (*Procyon lotor,* PRLO), eastern gray squirrel (*Sciurus carolinensis,* SCCA), and eastern cottontail rabbit (*Sylvilagus floridanus,* SYFL). As some individuals are unnecessarily taken from their habitat, there is a benefit to characterizing where they are collected. This study utilized Geographic Information System to examine the spatial relationship between the environment and originating locations of orphans presented to the University of Illinois Wildlife Medical Clinic within Champaign County from 2015-2020 (99 Virginia opossums, 80 common raccoons, 441 eastern gray squirrels, and 602 eastern cottontails). Environmental factors evaluated included percent tree canopy, land cover classification, and distance to water. Overall, these species were frequently found in highly developed areas (p < 0.001), near water (p < 0.027), with a low percent canopy (p < 0.001). Our analysis identifies environments associated with greater human-wildlife interactions and opportunities for targeted educational outreach.

## Introduction

Human-wildlife interactions occur when both parties are in spatial and temporal proximity [1]. These interactions are increasing as cities and towns expand into nature, fragmenting wildlife habitats [2]. In the United States, both urban and rural residential areas have recently been rapidly increasing [3]. Expanding developments impact local and regional ecosystems [4] and can displace wildlife [5]. Conversely, some species, such as common raccoons (*P. lotor*), thrive in developed areas and are overabundant [6]. This study analysis identifies environments with greater human-wildlife interactions to create opportunities for targeted educational outreach about orphans.

Orphans make up the largest number of cases seen by rehabilitation facilities [7–8]. A recent study of admission records from a suburban wildlife rehabilitation facility in central Ohio showed that 7 of the 20 most frequently admitted species were mammals [7]. That report found that eastern cottontail rabbits (*S. floridanus*) accounted for a quarter of all of the cases [7]. Concurrently, healthy neonatal and juvenile animals are commonly misidentified as orphans [9]. Raising orphans in captivity can negatively affect their survival, as habituation to humans reduces an animal’s response to predators and other hazards [10–11]. Tribe et al. found that some species displayed abnormal den and foraging behavior after release [10]. A survey of mammal wildlife rehabilitation centers indicated that rehabilitated animals had difficulty avoiding predators and some exhibited stereotypic behaviors compared to their wild counterparts [12]. Further, there is a benefit to reducing the intake of healthy orphans, allowing funds to be directed to animals that require more specialized care [13–14]. Understanding which environments are associated with greater human-wildlife interactions can help target outreach to reduce the intake of healthy orphans to rehabilitation centers.

This project focused on the four most common orphaned mammal species presented to a local wildlife rehabilitation center in Champaign County, Illinois, United States of America (US). The most common orphaned mammals presented were the Virginia opossum (*D. virginiana*), common raccoon (*P. lotor*), eastern gray squirrel (*S. carolinensis*), and eastern cottontail rabbit (*S. floridanus)*. Since the species studied are commonly found near human developments, this study hypothesized that all four species would be found in highly developed areas with a higher percentage of canopy cover that is close to water. Additionally, species-specific hypotheses were proposed about what types of environments each species would be commonly found in.

## Materials and Methods

This study identifies environments in Champaign County, Illinois, US, associated with greater human-wildlife interactions, measured by orphans found and brought in for care, for targeted educational outreach. This study utilized Geographic Information System to examine the spatial relationship between the environment and originating locations of orphans presented to the University of Illinois Wildlife Medical Clinic (WMC) from Champaign County from 2015 to 2020.

The most abundant species of orphaned mammals brought to the WMC were identified using RaptorMed (RaptorMed.com LLC, 2011). RaptorMed is an electronic medical record software utilized by wildlife rehabilitators and zoological collections. A comprehensive list of the mammals admitted with “orphaned” denoted on the animal’s problem list was generated. The four most common mammal species from that data, and therefore the species included in this study, were the Virginia opossum (*D. virginiana*), common raccoon (*P. lotor*), eastern gray squirrel (*S. carolinensis)*, and eastern cottontail rabbit (*S. floridanus*). A total of 1,222 animals (99 Virginia opossums, 80 common raccoons, 441 eastern gray squirrels, and 602 eastern cottontails) met the inclusion criteria for this study.

An advanced search of RaptorMed was utilized to create a list of cases admitted between 2015 to 2020. This date range was chosen because it contained more detailed records. Pertinent data (patient ID, admission date, species, age at intake, status, the reason for admission, and found address including city, county, state, and zip code) was exported into an Excel file (Microsoft Excel, 2019). Results were limited to the four mammal species of interest. Only subadult animals were included. Cases presenting from outside of Champaign County and those with no associated found address were excluded. Duplicate entries from animals admitted as a group were deleted. The final RaptorMed dataset contained 1,222 unique presentation records.

RaptorMed found address data was converted to latitude and longitude coordinates using an online geocoding platform (Dotsquare, 2021). The software generated an accuracy rating for each converted address. Scores ranged from 0 to 1, with 1 being the most accurate. All addresses with a rating under 1 were flagged and manually corrected. A second run of the data ensured proper conversion before spatial analysis.

Case distribution in Champaign County was visually represented using ArcGIS Pro (version 2.8, Esri, 2021). Layers represented the percent tree canopy, land cover classification, and distance to the nearest water source in the county. Data was extracted from ArcGIS detailing how the found location of each case correlated to measurements of percent canopy, land cover class, and distance to water on those respective layers. Results were exported onto an Excel file and separated by species.

Environmental factors evaluated included percent canopy, land cover class, and distance to water. The study utilized land cover classes from the National Land Cover Database (NLCD). NLCD data is generated by the United States Geological Survey (USGS) from satellite imagery using a complex modeling process. Each environmental category was split into high and low groups for statistical analysis.

High canopy cover was defined as 30.1% and above [14]. Landover classes 21-24 (Developed-Open Space, Developed-Low Intensity, Developed-Medium Intensity, and Developed-High Intensity) were defined as high development. Classes were separated into high development (21-24), which represented all developed areas, and low development (43, 81,82), which represented forest, hay/pasture, and crops. Land cover classes 43, 81, and 82 (Mixed Forest, Hay/Pasture, and Cultivated Crops) were defined as low development. According to Iverson, 79% of the forests in Illinois are within 300 m of streams [15]. The spatial resolution of the NLCD gridded data for water sources is 30 meters, which is not fine enough to pick up most streams. Distance to water was calculated in the GIS from high-resolution water feature data provided by the Champaign County GIS Consortium. Distances to water in this survey ranged from 0 to 2710 m and the median was 347.53 m. Low distance to water was defined as 0-347.53 m and high distance to water was greater than 347.53 m. Tables were created dividing the cases into these sections for statistical analysis.

Stata software (version 16.1, 2020, StataCorp LLC) was used to run a two-sample test of proportions. The results from this test indicate whether the high and low categories within one species differed. The null hypothesis (H0) for each run was: p1 = p2. The alternative hypothesis was (Ha): p1 ≠ p2. If the Z test was significant, the null hypothesis was rejected, indicating a significant difference between the two groups.

## Results

The four species studied were frequently found by members of the public in the Champaign-Urbana, IL area (Fig 1) in highly developed areas (p < 0.001) (Table 1, Fig 2, Fig 3), with a low percent canopy (p < 0.001) (Table 2, Fig 4), that are close to water (p < 0.027) (Table 3, Fig 5). These findings are summarized in a single image (Fig 6).

**Figure 1.**
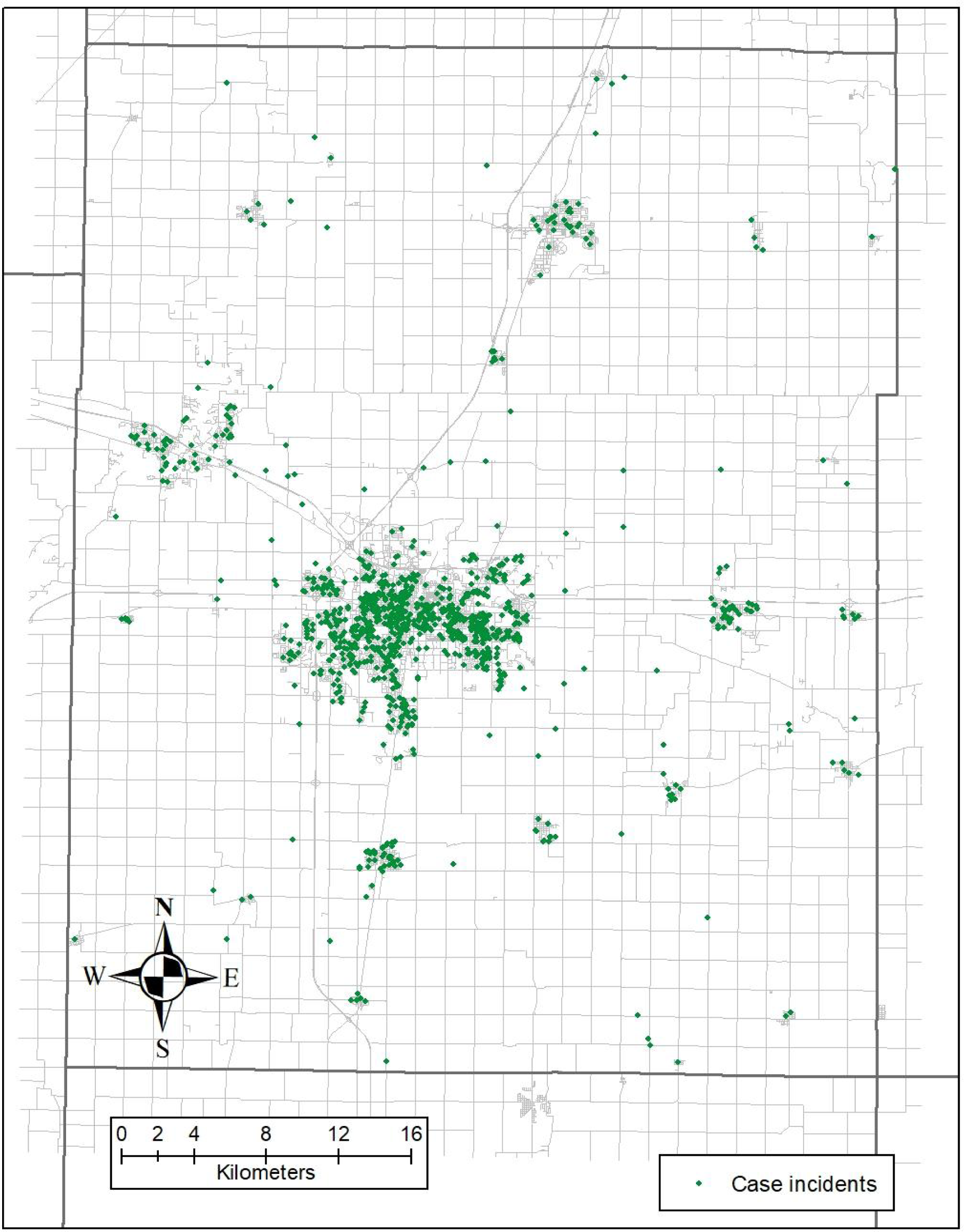
County Case Distribution. Case distribution (⬥) within Champaign County, IL US. The majority of cases cluster around the Champaign-Urbana, IL area.

**Table 1.**
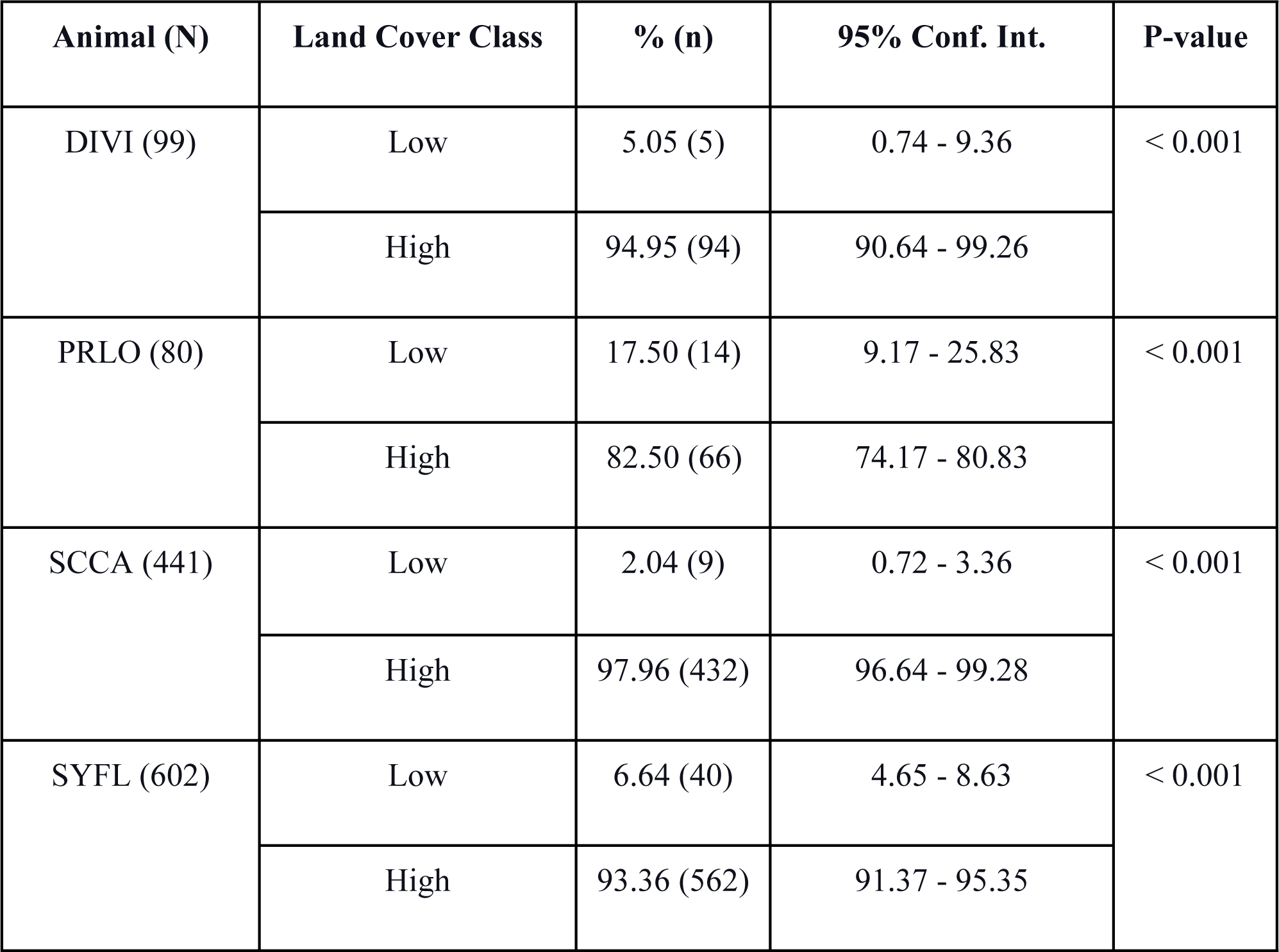
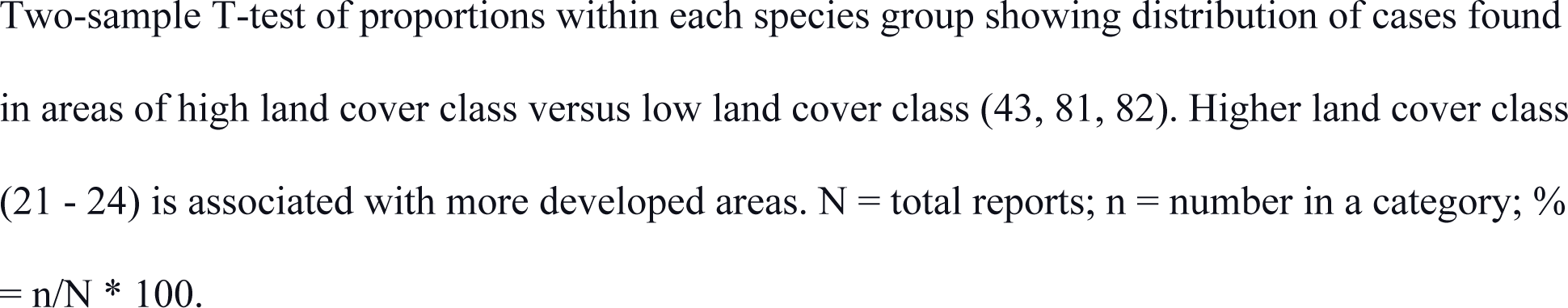
Land Cover Class Table.

**Figure 2.**
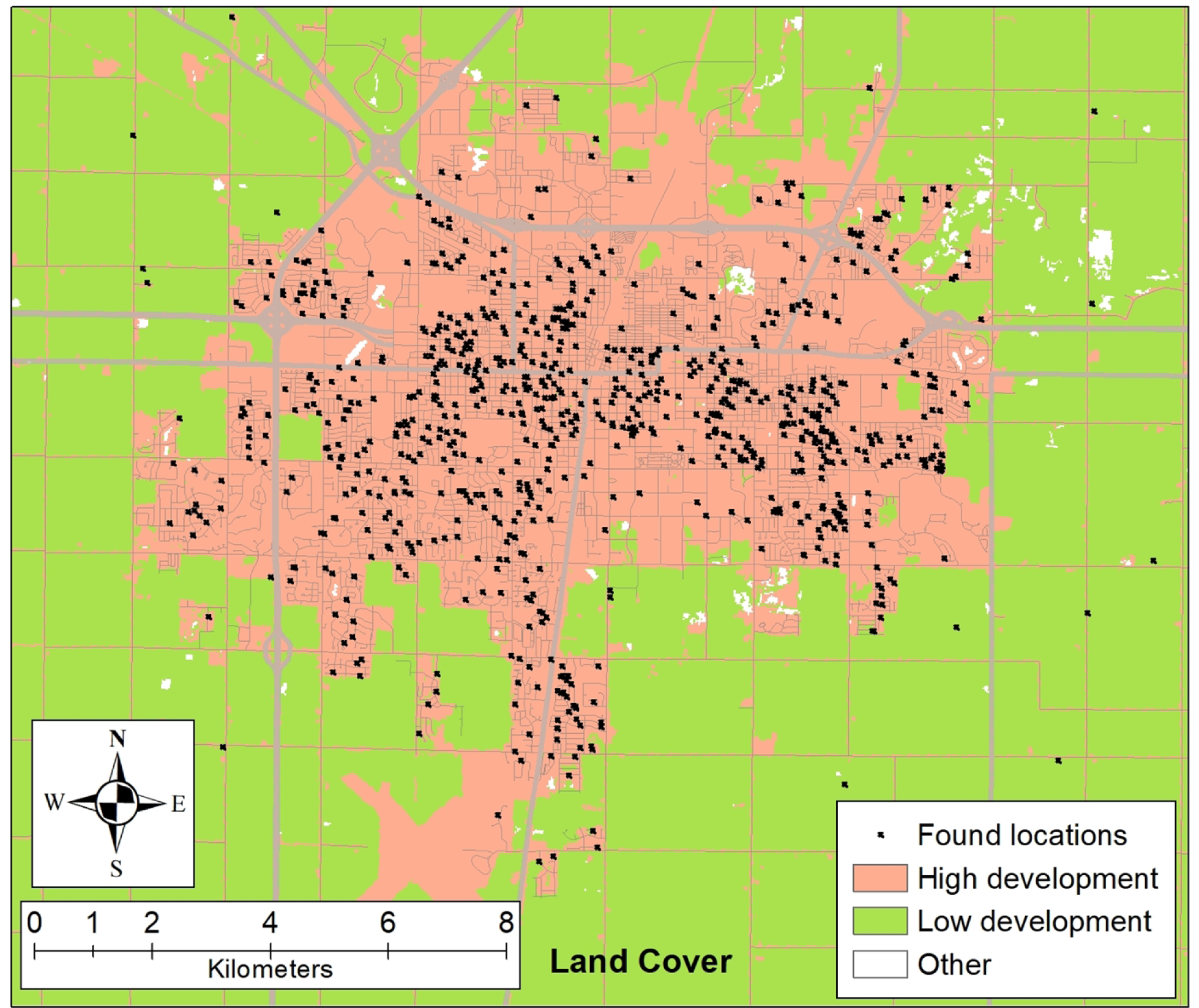
Development Case Distribution. Case distribution on the map layer of the land cover class showing whether cases were found in areas of low or high development.

**Figure 3.**
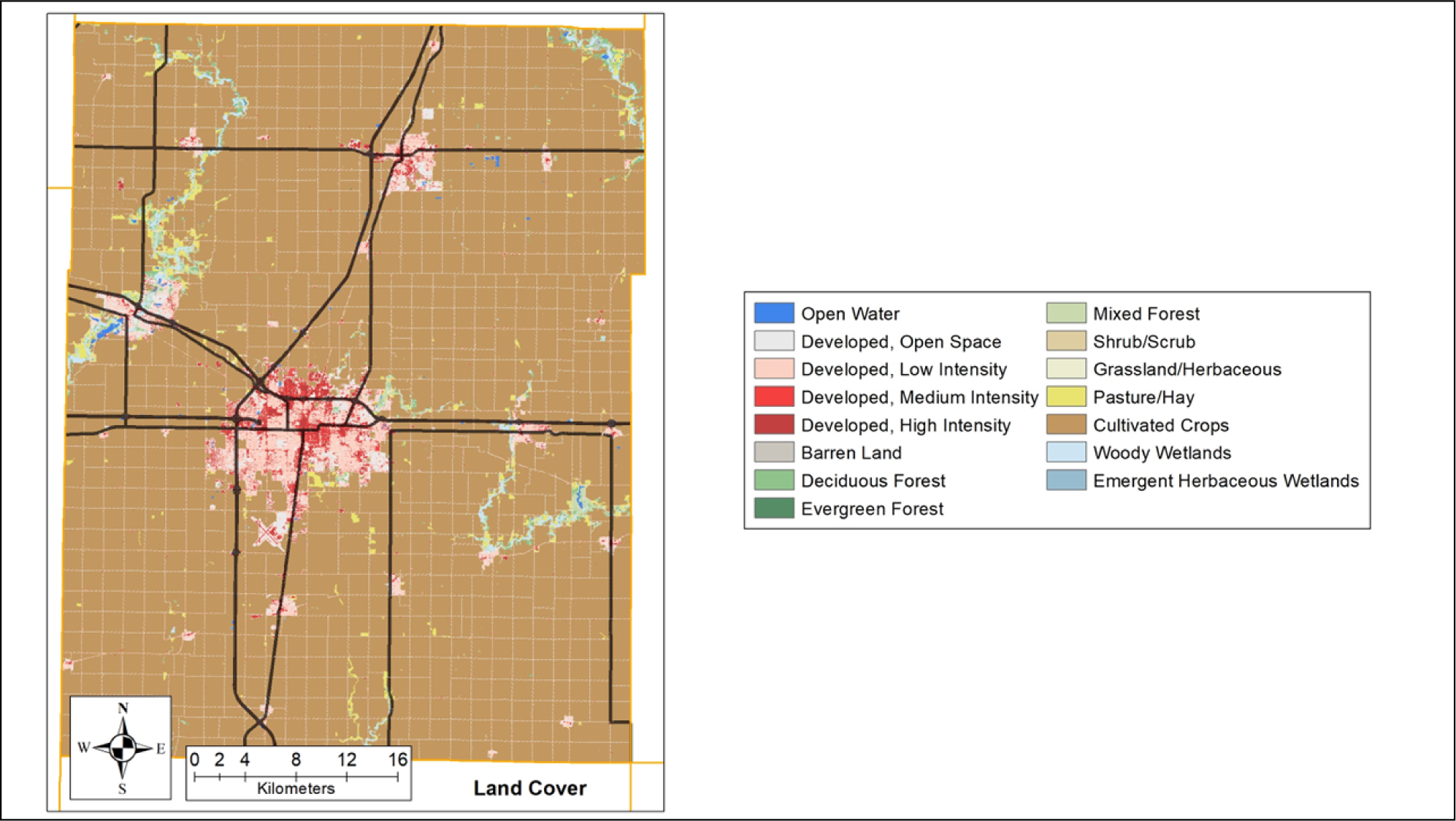
Land Cover Class Case Distribution. Case distribution (x) on the map layer of land cover class in Champaign County, IL. The color of the background represents the land cover class type.

**Table 2.** Percent Canopy Table.

| Animal (N) | Percent Canopy | % (n) | 95% Conf. Int. | P-value |
| --- | --- | --- | --- | --- |
| DIVI (99) | Low | 89.90 (89) | 83.96 - 95.83 | < 0.001 |
|  | High | 10.10 (10) | 4.17 - 16.04 |  |
| PRLO (80) | Low | 92.50 (74) | 86.73 - 98.27 | < 0.001 |
|  | High | 7.50 (6) | 1.73 - 13.27 |  |
| SCCA (441) | Low | 96.15 (424) | 94.35 - 97.94 | < 0.001 |
|  | High | 3.85 (17) | 2.06 - 5.65 |  |
| SYFL (602) | Low | 95.51 (575) | 93.86 - 97.17 | < 0.001 |
|  | High | 4.49 (27) | 88.69 - 93.37 |  |

Two-sample T-test of proportions within each species group showing distribution of cases found in areas of high percent canopy (> 30%) versus low percent canopy (0 - 30%). N = total reports; n = number in a category; % = n/N \* 100.

**Figure 4.**
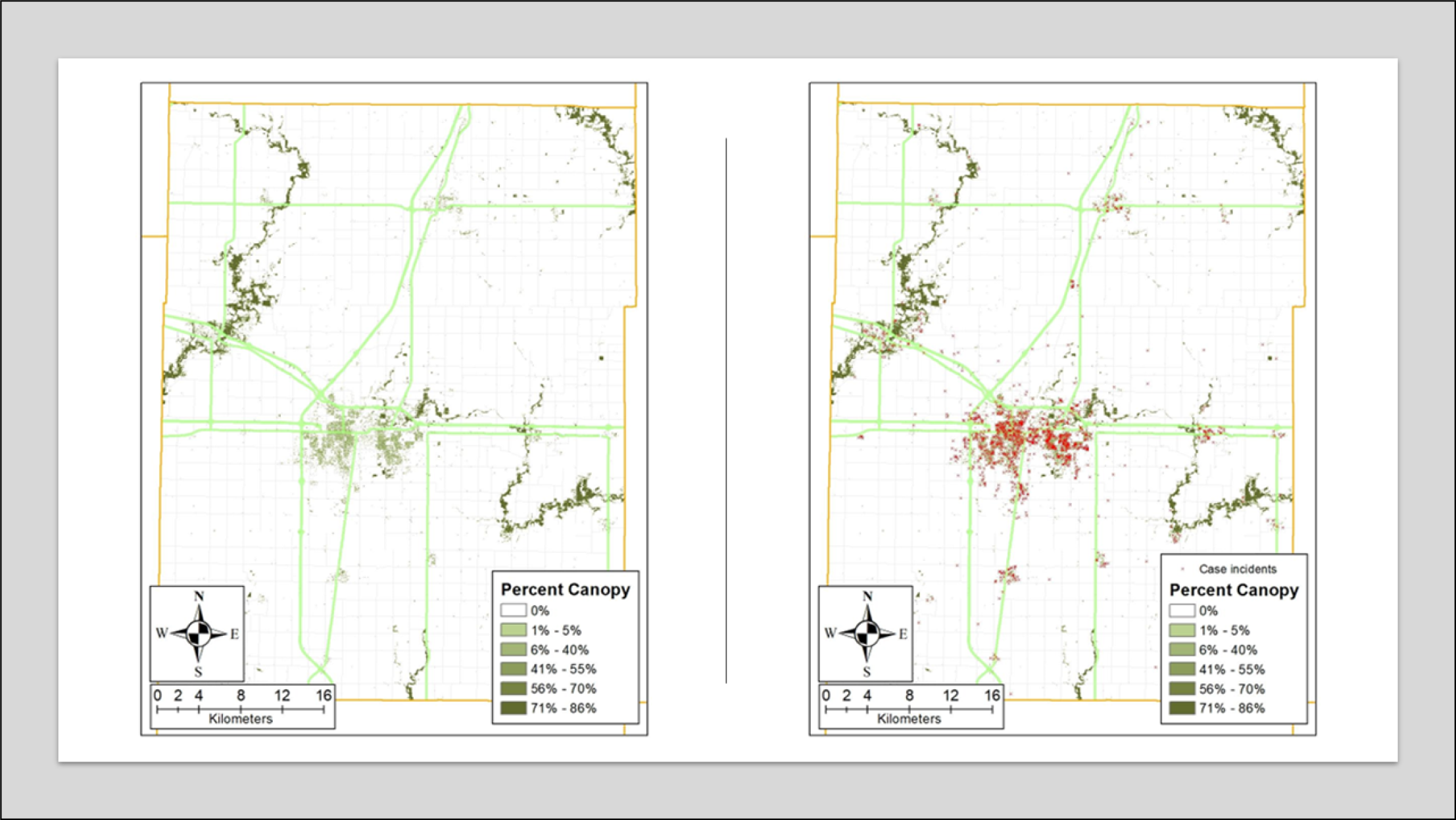
Percent Canopy Case Distribution. Case distribution (x) on the map layer of percent canopy in Champaign County, IL.

**Table 3.** Distance to Water Table.

| Animal (N) | Distance to Water<br>(meters) | % (n) | 95% Conf. Int. | P-value |
| --- | --- | --- | --- | --- |
| DIVI (99) | Low | 53.54 (53) | 43.71 - 63.36 | 0.32 |
|  | High | 46.46 (46) | 36.64 - 56.29 |  |
| PRLO (80) | Low | 58.75 (47) | 47.96 - 69.54 | 0.027 |
|  | High | 41.25 (33) | 30.46 - 52.04 |  |
| SCCA (441) | Low | 41.50 (183) | 36.90 - 46.10 | < 0.001 |
|  | High | 58.50 (258) | 53.90 - 63.10 |  |
| SYFL (602) | Low | 54.49 (328) | 50.51 - 58.46 | 0.002 |
|  | High | 45.51 (274) | 41.54 - 49.49 |  |

Two-sample T-test of proportions within each species group showing distribution of cases found in areas with a higher distance to water (> 347.53 m) versus areas with a low distance to water (0 - 347.53 m). N = total reports; n = number in a category; % = $n/N * 100$

**Figure 5.**
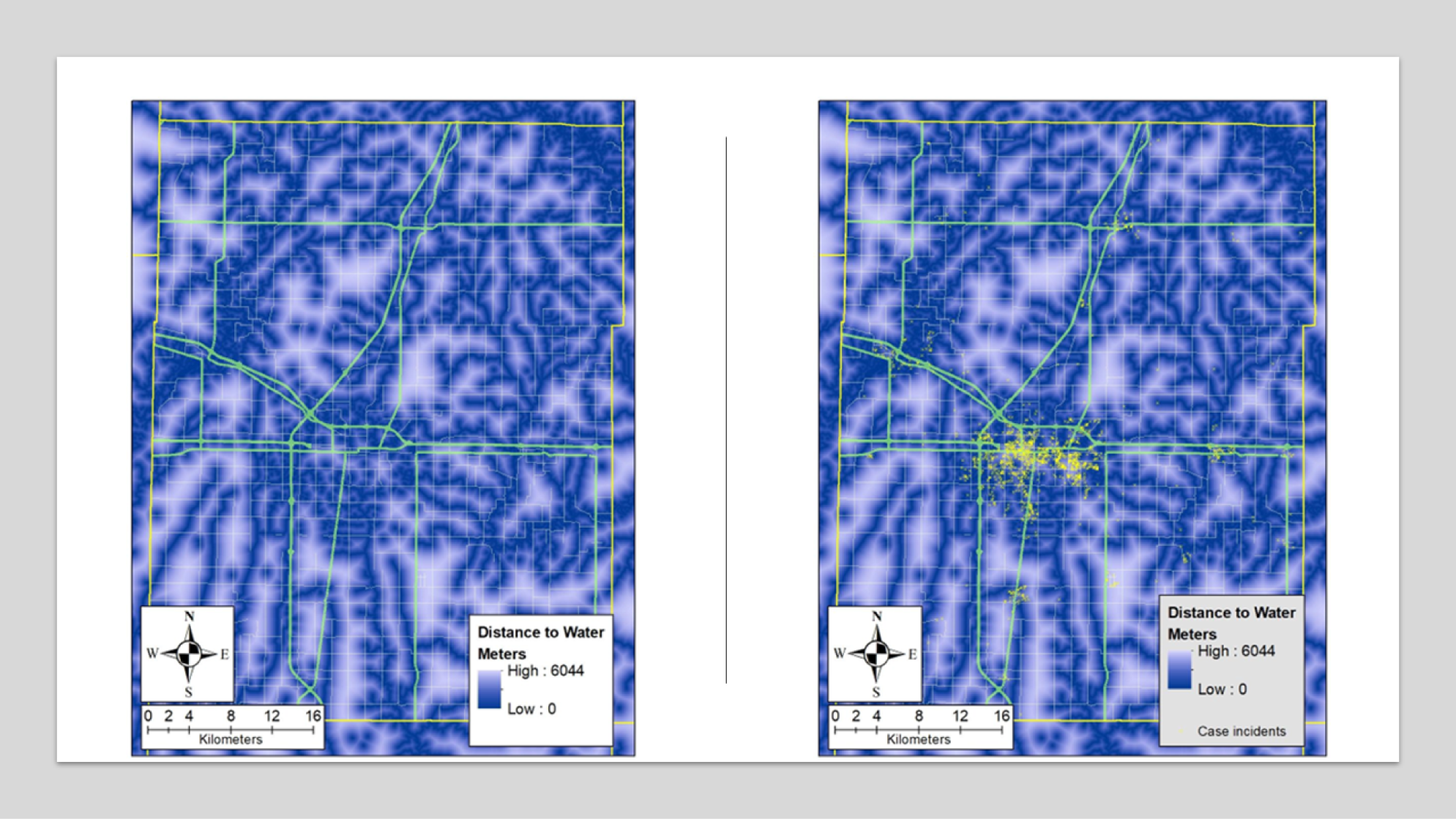
Distance to Water Case Distribution. Case distribution (x) on the map layer of distance to water in Champaign County, IL.

**Figure 6.**
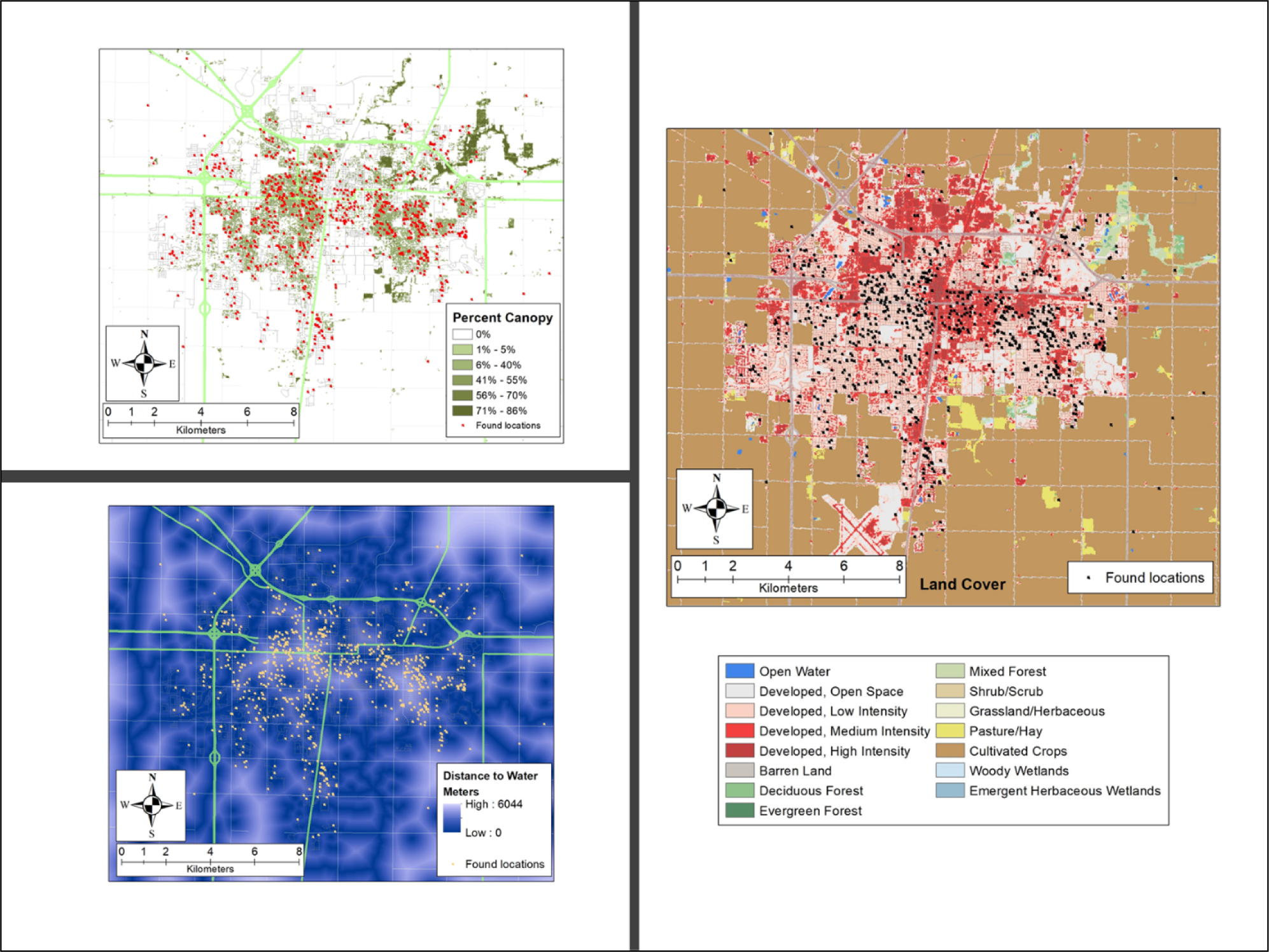
Case Distribution with Environmental Factors. Case distribution on the map layers of percent canopy (top), land cover class (middle), and distance to water (bottom) in the Champaign-Urbana, IL area.

Virginia opossums and common raccoons were frequently found in areas of high development Table 1 (p < 0.001). However, both were found in areas of low percent canopy Table 2 (p < 0.001). Common raccoons were most found close to water sources (Table 3). There was no significant difference in distance to water for Virginia opossums (Table 3). Eastern gray squirrels were found furthest from water sources compared to other species (Table 3). This species favored highly developed areas with low canopy cover Tables 1 and 2. Eastern cottontails were found in areas with low percent canopy (Table 2) that were highly developed (Table 1). While it was hypothesized that eastern cottontails would be found the furthest from water, the data from this study showed this species was more likely to be found near water (Table 3).

## Discussion

This study identified environments associated with more human-wildlife interactions in Champaign County, Illinois, US. The scope of the project focused on the four most common orphaned mammal species presented to a local wildlife rehabilitation center. It was hypothesized that the four types of orphans would be found in highly developed areas, close to water, and in more forested areas. The results showed that these species were found in highly developed areas, close to water, with a low percent canopy T. In Champaign County, these environmental characteristics are found in urban areas. Most cases originated from the Champaign-Urbana, IL area, which is also where the rehabilitation facility is located. While this project focused on wildlife presenting from Champaign County, IL, a similar approach can be applied to other regions.

Similar to a previous study from the American Midwest [7], this study found that more animals were admitted from urban and suburban areas. In another study, more human-wildlife interactions in northern New York clustered around suburban and low-density residential areas [17]. The results in this study parallel these findings, as the majority of cases came from highly developed areas.

Species that respond well to urbanization are typically generalists with broad dietary and habitat requirements. Mesopredators, such as the Virginia opossum *(D. virginiana)* and common raccoon *(P. lotor)*, are generalists who respond well to human developments and habitat fragmentation [18, 6]. Furthermore, these species have similar dens, habitats, and diets and may have overlapping ranges [19]. Both species prefer wooded areas in their natural habitats [6]. Based on the literature, it was hypothesized that common raccoon *(P. lotor)* and Virginia opossum *(D. virginiana)* orphans would be found in highly developed areas with high tree cover that are close to water. Further, it was hypothesized that raccoons *(P. lotor)* would be found in the most developed areas compared to the three other species. Raccoons *(P. lotor)* exploit anthropogenic resources more efficiently than other mesocarnivores [6]. Urban raccoons *(P. lotor)* have increased survival, annual recruitment, and site fidelity, and these factors may have led to their abundance [20]

Conversely, it was hypothesized that more eastern gray squirrels (*S. carolinensis*) would be found in less developed, forested areas that were close to water. Gray squirrels (*S. carolinensis*) are commonly found in densely wooded [21, p. 217] and urban areas [21, p. 221]. In central and northern Illinois, an area must be at least 20% forest to support a gray squirrel population (*S. carolinensis*) [22, p. 215]. Peplinski & Brown surveyed faculty experts at 536 Canadian and American college campuses and found that 63% of campuses had eastern gray squirrels, and eastern gray squirrels and fox squirrels shared similar resources on college campuses [23]. The two species coexisted in the same habitats by partitioning resources in their native and introduced ranges [23]. Eastern grey squirrels (*S. carolinensis*) were abundant in areas of high human development and low canopy covers such as urban parks, forests, and neighborhood habitats, and were nearly absent in cemeteries and golf courses [24]. Our results are similar because all of the species studied were found in highly anthropogenic areas.

Further, it was hypothesized that more eastern cottontails (*S. floridanus*) would be found in less developed, forested areas that are further from water. This hypothesis was based on the type of habitats they prefer. In central Illinois, rabbits (*S. floridanus*) are locally abundant in diverse, patchy areas with moderate amounts of crops, grassland, and at the edges of woods [25]. Paths and roads are also important elements of their ranges [21]. Due to this natural history, it was expected this species would be found in less developed areas with less tree cover. Rabbits (*S. floridanus*) are also able to successfully inhabit parks in urban environments [26]. Annual survivorship was similar to those in natural environments, however, urban rabbits had smaller ranges [26].

In this study, the common raccoon *(P. lotor)* was found the closest to water sources; this finding is supported by their natural history. Raccoons *(P. lotor)* are found almost anywhere water is located [21]. Common raccoon tree dens averaged 23-33 m from water sources and ground burrows averaged 75-101 m from water in west-central Illinois [27].

Illinois is mostly covered by agricultural land [27, 3]. As a result, only a small portion of Illinois is considered high-quality for wildlife [29]. Compared to the total land area, agricultural land occupies 71.5%, and developed areas occupy 11.5% of Illinois [28]. Natural areas occupy the remaining portion and 15.6% of the state is forested [28]. Historically, Illinois’ forests have been located near water since streams are an effective fire barrier [16]. Consequently, Illinois’ forests are highly fragmented; which negatively impacts species in need of continuous habitats [16].

Proximity to water is one of the most important factors in how vertebrates utilize space [30]. Illinois has over 21,200 km of streams and rivers [31, p.154]. In Central Illinois, 79% of forests are within 300 meters of streams, and 22% of forests are within 30 meters of streams [16]. This pattern occurs due to the difficulty of growing crops on the slopes of the streams [16]. The Middle Fork, Salt Fork, and Windfall Creek streams had the highest quality wetlands and nearby natural areas for wildlife in central Illinois [32].

Tree canopy cover is a proxy used to assess an area’s forests [33]. Tree cover has been declining by about 36 million trees per year in urban and community areas nationwide [33]. The greatest loss was in urban areas (-1%), coinciding with a 1% increase in grass and impervious cover [33]. Community areas had a 0.7% loss of tree cover with a 0.6% increase in impervious cover [33]. Tree canopy cover averages 15.6% statewide (SD = 1.1%), 26.4% (SD = 2.4%) in urban areas, and 14.7% (SD= 1.2%) in rural areas in Illinois [34]. This evaluation was performed using trained evaluators who interpreted Google Earth^Ⓡ^ images. A recent evaluation performed utilized Google Street Corridor to evaluate the percent canopy and establish low, medium, and high categories [15]. The low cover was classified as 0-15.0%, medium cover as 15.1-30.0%, and high cover as 30.1-62.0% [15].

A major limitation of this study is that intakes to the wildlife rehabilitation facility are limited by what animals are found and presented by the public. Certain mammal species that are abundant in developed areas, such as juvenile foxes and coyotes, may be less likely to be handled by members of the public due to safety concerns. The results of this study are also limited by the finder’s ability to report the found locations of the animals presented for care. Intakes without a found address were excluded, which could bias the conclusions drawn. It can further be suspected that socioeconomic factors, such as an area’s family income and education level, could impact animal admissions. A recent meta-analysis showed that urban dwellers had the most positive perception of wildlife [35]. Despite these limitations, the results of this study demonstrate the value of using spatial analysis to identify areas of increased human-wildlife interactions.

The results of this study can help wildlife rehabilitation and conservation centers target areas for education and fundraising. This study explored the relationship between where wildlife were found with land cover class, percent canopy cover, and distance to water. Based on these results and other literature investigating these relationships, expanding urban developments may create more human-wildlife interactions with generalist animal species. These interactions lead to more chances for the public to find orphaned wildlife. Future studies can use the same approach to study the remaining Illinois counties and other states.

## Acknowledgments

The authors would like to thank the University of Illinois Wildlife Medical Clinic, the University of Illinois at Urbana-Champaign College of Veterinary Clinical Medicine, and the University of Illinois at Urbana-Champaign Department of Pathobiology for their contributions to this project.

